# NACraft: Programmatic nucleic-acid aptamer design via all-atom structure-model feedback

**DOI:** 10.64898/2026.08.15.744087

**Authors:** Heqin Zhu, Jiaqi Wang, Weibo Zhao, Yuzhi Xu, Huang Su, Jianmin Wang, Qinghan Wang, Yuntao Yu, Ziyi You, Gang Du, Pheng Ann Heng, Liqin Zhang, Odin Zhang

## Abstract

Protein–nucleic-acid interactions underpin diverse biological processes and provide a basis for molecular sensing, regulation and therapeutic intervention. However, the coupled dependence of aptamer function on nucleotide sequence, three-dimensional folding and target binding makes rational RNA and DNA binder design challenging. Here we present NACraft, a training-free and programmatic framework for all-atom nucleic-acid aptamer design based on backpropagation through structure-model feedback. By composing binding, sequence-similarity and anti-binding constraints, NACraft supports de novo generation, similarity-guided sampling and target-selective design within a unified optimization framework, without task-specific training or fine-tuning. Computational experiments showed that NACraft generated high-confidence candidates de novo across diverse protein targets, with further improvements achieved through similarity-guided design for both RNA and DNA complexes. Its target-selective design capability was further validated in silico, with 69.44% of paired candidates generated to favour the positive target EGFR over the off-target HER2. Under matched independent AlphaFold3 evaluation, NACraft achieved better performance than ODesign in 10 of 11 NA-12 targets and 17 of 20 protein target–length settings. Together, these results demonstrate the effectiveness and versatility of NACraft and extend structure-model hallucination toward programmatic nucleic-acid aptamer design.

**Code:** https://github.com/OTEAM-AI4S/NACraft

## 1 Introduction

Nucleic-acid aptamers are short RNA or DNA molecules that demonstrate canonical and non-canonical base pairings [54, 55], and further fold into three-dimensional structures [38] that bind molecular targets with high affinity and specificity [15]. Their compact size, synthetic accessibility and chemical programmability have enabled applications in biosensing [46], diagnostics [41], targeted delivery [42], molecular regulation [50] and therapeutic development [6]. Most functional aptamers are discovered through iterative selection and enrichment, followed by optimization of experimentally identified sequence families [16]. These approaches have produced useful binders, yet leave a central inverse-design problem unresolved: given the structure of a target, how can one construct a previously unknown nucleotide sequence that folds into a complementary binding surface? This problem is challenging because nucleotide identity, base-pairing, tertiary folding and intermolecular binding are strongly coupled.

Computational molecular design is increasingly shifting from candidate screening towards direct generation. Classical nucleic-acid design methods optimize sequences against predefined secondary-structure ensembles and are effective when the desired base-pairing pattern is known [11, 49]. Selection-informed approaches provide an alternative route by learning directly from experimentally enriched sequences. For example, InstructNA [52] combines high-throughput SELEX with nucleic-acid large language models (NA-LLMs) [20, 31, 53] to generate functional nucleic acids without requiring target three-dimensional structures. More recently, diffusion-based generative models and sequence–structure co-design models [7, 13, 14, 27, 37, 39, 43] have enabled joint generation of molecular sequences and structures. In the nucleic-acid setting, however, these approaches have largely focused on generating individual RNA molecules, without explicitly conditioning design on a binding target, and their applicability to DNA aptamer design remains limited. ODesign [51] partially addresses these limitations through an all-to-all generative framework that supports protein-binding RNA and DNA design within a unified molecular representation. Nevertheless, its current formulation is centred primarily on de novo generation and provides limited support for design settings that arise frequently in practical aptamer engineering, such as improving an existing functional aptamer or explicitly optimizing selectivity between competing targets.

Recent advances in structure-prediction models have opened a complementary route to molecular design through structure-model hallucination [9, 21, 34, 35]. Rather than sampling from a separately trained generative model, hallucination treats the sequence itself as an optimizable variable and backpropagates structural objectives through a pretrained folding model, thereby enabling new design tasks to be specified at inference time without task-specific training or fine-tuning. This strategy has so far been developed predominantly for proteins. BindCraft [34] uses AlphaFold-based optimization [22] for de novo protein-binder design; Protein Hunter [35] combines hallucination with diffusion-based structure prediction and iterative redesign; BoltzDesign1 [9] inverts an all-atom structure predictor [44] for generalized protein-binder design; and SwitchCraft [21] introduces compositional objectives for programmatic multistate protein design. Although some of these frameworks can include nucleic acids in the molecular context, the optimized molecule remains a protein rather than an RNA or DNA aptamer. Extending hallucination to nucleic-acid design therefore requires a formulation that operates directly over nucleotide and supports the broader range of objectives encountered in aptamer engineering. In particular, a general framework should accommodate three complementary settings: *de novo design*, which explores novel aptamer sequence space; *similarity-guided design*, which leverages existing aptamer sequences or motifs to guide improved designs; and *target-selective design*, which optimizes differential binding between desired and competing targets to favour target binding while reducing off-target interactions.

To address this gap, we introduce NACraft, a training-free, programmatic and all-atom hallucination framework developed specifically for RNA and DNA aptamer design. NACraft represents an aptamer as optimizable nucleotide logits constrained to the corresponding RNA or DNA nucleotide space. It adapts structure-model inversion to nucleic-acid representations and compositional objectives for target binding, intramolecular contacts, sequence guidance and target selectivity, enabling distinct aptamer-design tasks to be specified within a unified optimization framework. By changing the molecular contexts and objective terms, the same optimization framework supports de novo, similarity-guided and target-selective design without retraining the underlying structure model or constructing separate task-specific pipelines. NACraft organizes this process into four stages: design-context statement, differentiable sequence optimization, NA-MPNN-based refinement and diversification [3], and independent all-atom validation and candidate filtering with AlphaFold3 (AF3) [1]. This formulation provides a unified interface for translating user-defined target-binding objectives into candidate RNA or DNA aptamers.

We systematically evaluated NACraft framework across multiple design settings in silico. First, we performed de novo RNA design against five therapeutically relevant protein targets. The resulting candidates reached maximum AF3 ipTM values of 0.63–0.88, with approximately 20% exceeding an ipTM threshold of 0.6 on average, demonstrating the capacity of NACraft for de novo aptamer generation. Next, we evaluated NACraft on NA-12, a benchmark comprising 12 structurally diverse nucleoprotein complexes from the PDB [4]. Across this benchmark, similarity-guided design consistently improved ipTM, pLDDT, and iPAE over de novo design, yielding high-confidence binding predictions and structural models for both RNA–protein and DNA–protein complexes. We then assessed target-selective design using EGFR [25] as the positive target and HER2 [8] as the negative target. Overall, 69.44% generated candidates favoured EGFR while disfavouring HER2, demonstrating the ability of NACraft to integrate positive- and negative-target objectives within a single optimization framework. Furthermore, we compared NACraft with the generative model ODesign [51], where NACraft achieved a higher target-level median ipTM for 10 of 11 NA-12 targets and in 17 of 20 therapeutically relevant protein target–length settings, demonstrating the superiority of NACraft for aptamer design. Finally, we performed ablation studies to quantify the contributions of similarity-loss weighting, the redesign procedure, broader sequence sampling and different optimization stages. Together, these results establish NACraft as a general framework for training-free, programmatic and all-atom protein binder design.

## 2 Results

### 2.1 Overview of NACraft for unified aptamer design

NACraft formulates RNA and DNA aptamer generation as programmatic sequence optimization under all-atom structure-model feedback (Fig. 1). The framework supports three complementary design modes (Fig. 1a). 1) In de novo protein-binding design, NACraft starts without a reference aptamer and searches for nucleotide sequences predicted to bind a specified target. 2) In similarity-guided design, a known or putative aptamer provides a soft similarity constraint while the interface remains free to remodel. 3) In target-selective design, candidate sequences are optimized to favour binding to the desired target over competing or off-target molecules. These modes are implemented through a unified optimization engine by changing the molecular contexts and compositional objective terms, rather than retraining the underlying structure model.

**Figure 1.**
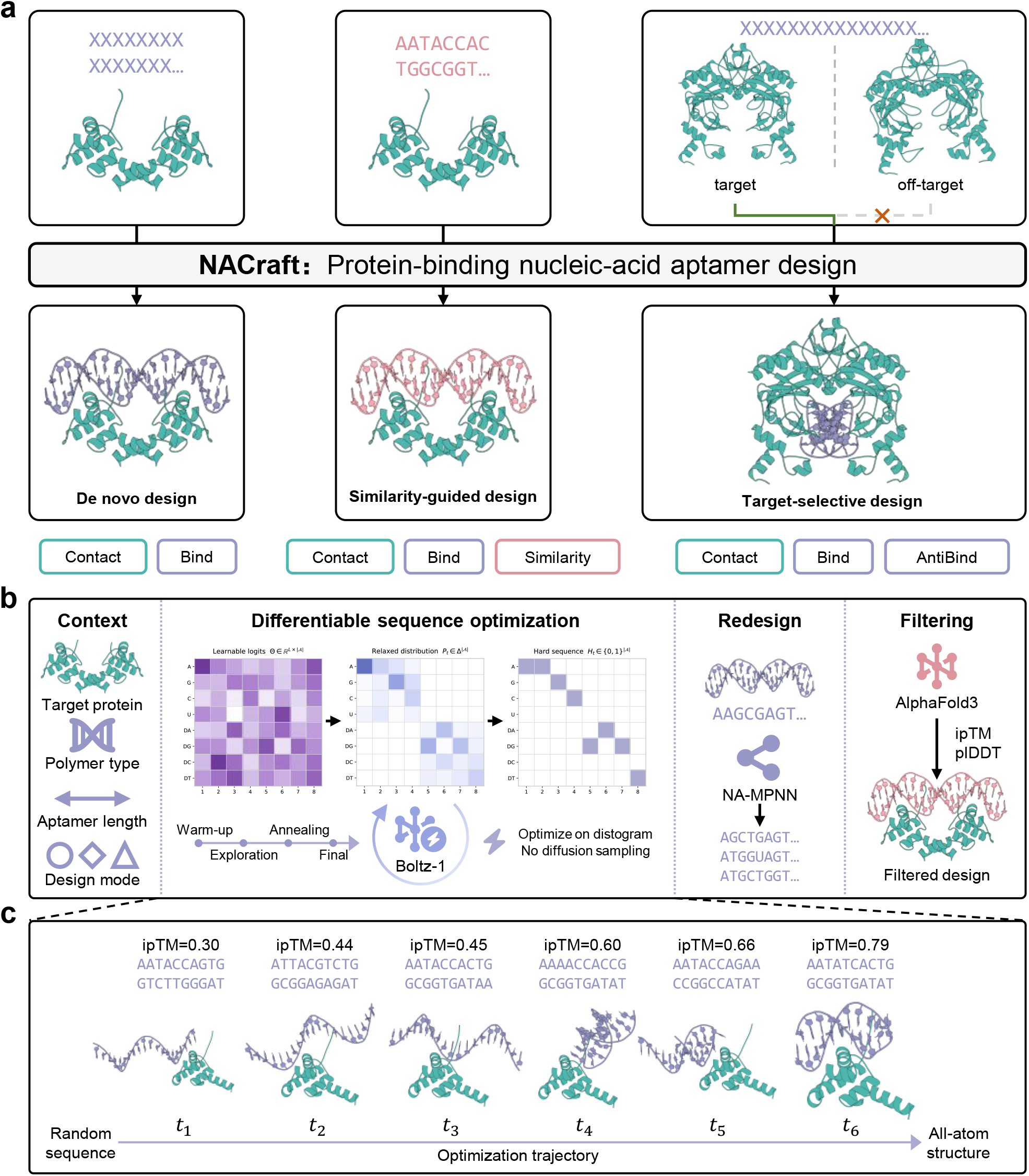
Overview of NACraft for programmatic aptamer design. **a**, Three aptamer design modes: de novo protein-binding design, similarity-guided remodeling and target-selective design against positive and negative targets. **b**, Four-stage NACraft workflow comprising design-context statement, differentiable sequence optimization through Boltz-1 distogram feedback, NA-MPNN sequence refinement and diversification, and independent AF3 validation and candidate filtering. **c**, Optimization trajectory from relaxed nucleotide logits to a discrete aptamer sequence through warm-up, exploration, annealing and final optimization phases, followed by all-atom structure validation.

The NACraft workflow comprises four stages (Fig. 1b). First, the design context specifies the protein targets, nucleic-acid classes, aptamer length and objective terms. Second, NACraft performs differentiable sequence optimization through Boltz-1 [44] structure-model feedback. Within this inner loop, relaxed nucleotide logits are progressively converted into a discrete sequence through warm-up, exploration, annealing and final low-temperature phases (Fig. 1c). Geometric and interaction losses are evaluated from Boltz-1 distogram outputs and backpropagated to the logits without diffusion-based coordinate sampling. Third, the optimized sequences are materialized as all-atom structures and refined and diversified using NA-MPNN. Fourth, all candidates undergo independent AF3 prediction and confidence-based filtering. Separating differentiable search, structure-based diversification and independent validation allows each stage to use a complementary model without task-specific retraining.

In the following sections, we first evaluate de novo aptamer generation across five therapeutically relevant protein targets and then use NA-12 to assess similarity-guided design across RNA and DNA aptamers. We next assess target selectivity using an EGFR-over-HER2 selective-design task. To benchmark overall design performance, we compare NACraft with the diffusion-based ODesign framework under matched AF3 rescoring. We then perform ablation analyses of sequence guidance, NA-MPNN refinement and optimization dynamics.

### 2.2 NACraft enables de novo aptamer design

A general de novo framework should generate candidate aptamers across distinct protein targets without relying on reference sequences. We evaluated this capability using B7-H3 [2], PD-L1 [36], CD3*δ* [12], TNFR1 [26] and FGFR2 [17] at aptamer lengths of 20, 30, 40 and 50 nucleotides (Fig. 2; Supplementary Tables 1 and 3). NACraft generated 300 sequences for each target–length setting, totalling 6,000 designs across five targets and four lengths, which were evaluated using AF3-derived ipTM and pLDDT. As shown in Fig. 2a and b, the best candidates reached ipTM values of 0.63, 0.80, 0.81, 0.85 and 0.88 for CD3*δ*, PD-L1, TNFR1, B7-H3 and FGFR2, respectively. Across these targets, ipTM decreased modestly with increasing aptamer length, although high-confidence candidates were obtained over a broad range of length settings. Besides high ipTM scores, the generated candidates also exhibited high interface pLDDT values, which were positively correlated with ipTM (Pearson *r* = 0.731; Fig. 2c), further supporting the quality of the generated interfaces. The redesign procedure further improved candidate quality (Fig. 2d). We next quantified both the fraction of candidates exceeding an AF3 ipTM threshold of 0.60 and their contact with designated hotspot residues (Fig. 2e). The fraction exceeding the ipTM threshold ranged from 1.08% for CD3*δ* to 41.50% for FGFR2, with values of 12.50%, 17.17% and 34.42% for B7-H3, PD-L1 and TNFR1, respectively. Hotspot contact, defined by an RNA–protein distance within 5 Å of the designated hotspot residues, ranged from 2.42% for CD3*δ* to 73.67% for TNFR1. Overall, NACraft generated high-confidence candidates against all five protein targets without a reference aptamer or target-specific training. The best candidates for each target–length setting are visualized in Fig. 2f using PyMOL [10].

**Figure 2.**
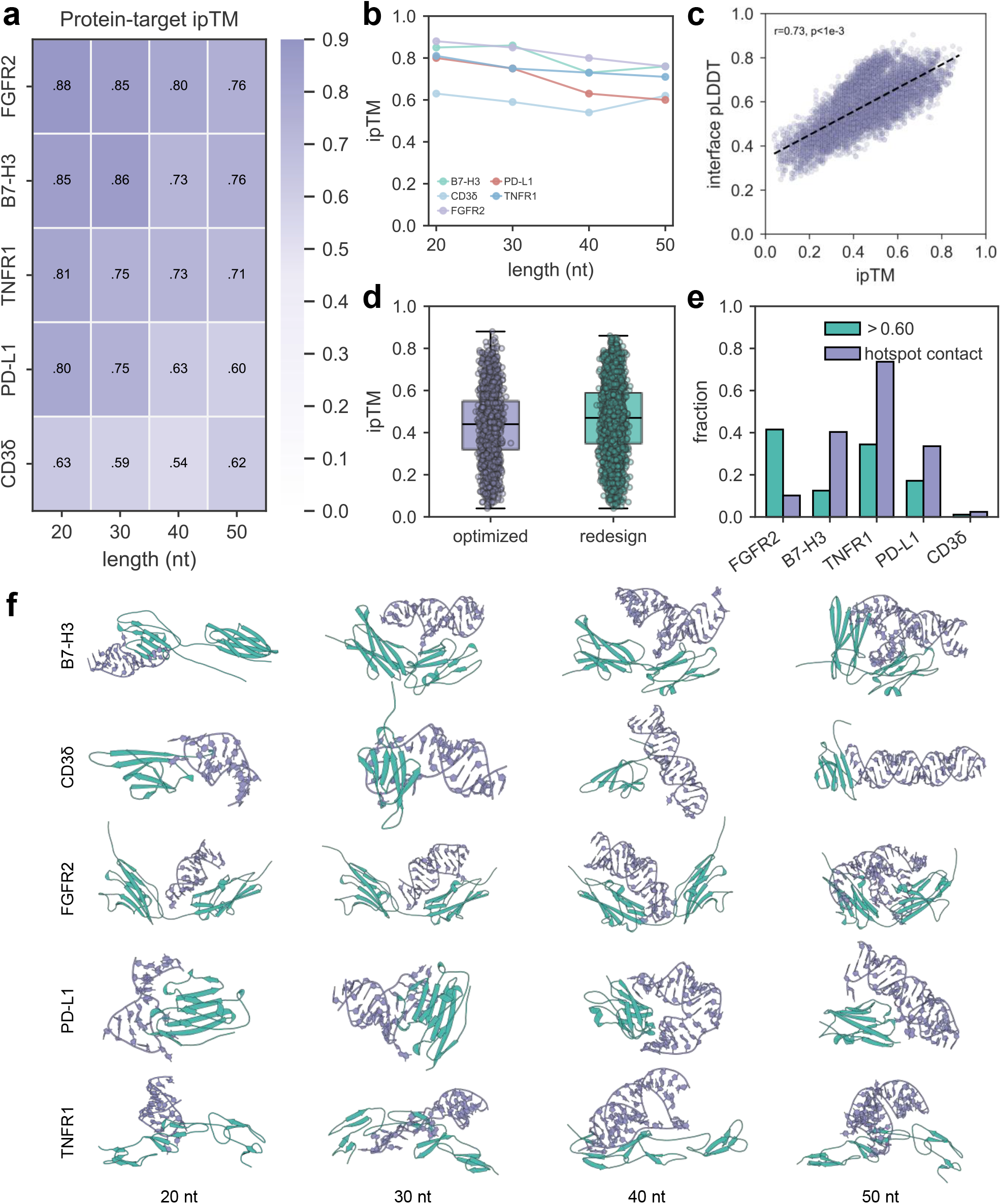
NACraft generated high-confidence RNA candidates against five therapeutically relevant protein targets. **a**, Target-by-length maximum AF3 ipTM across B7-H3, PD-L1, CD3*δ*, TNFR1 and FGFR2 at aptamer lengths of 20, 30, 40 and 50 nucleotides. **b**, Maximum AF3 ipTM across aptamer lengths for each protein target. **c**, Candidate-level AF3 ipTM versus normalized interface pLDDT across all therapeutically relevant protein target designs; the black dashed line denotes the linear fit, with the Pearson correlation reported in the panel. **d**, AF3 ipTM distributions for directly optimized parents and NA-MPNN-redesigned candidates. **e**, Fractions of candidates with AF3 ipTM greater than 0.60 and with predicted hotspot contact, shown separately for each protein target. **f**, Representative high-ipTM AF3-predicted complexes across the 20 protein–length settings, with proteins in green and RNA aptamers in purple. The analysis comprises 6,000 candidates across 20 protein–length settings.

### 2.3 NACraft incorporates sequence priors for similarity-guided aptamer design

Similarity-guided design addresses a complementary practical setting in which information from an existing aptamer should guide the search without preventing interface remodeling. To evaluate this capability across both nucleic-acid classes, we constructed NA-12 from six protein–RNA and six protein–DNA contexts (PDB IDs 9XZR [30], 8X0N [47], 8SWC [19], 9RVP [5], 9NY9 [24], 8SWB [18], 8VX4 [48], 8SLN [28], 8C58 [45], 7UXY [33], 8U0P [23] and 7UV7 [32]) and evaluated de novo and similarity-guided design under the same AF3 protocol (Supplementary Tables 1 and 2). Across these experiments, 7,200 AF3-scored candidates were evaluated, spanning both de novo and similarity-guided design across all 12 targets (structures are visualized in Fig. 3 using ChimeraX [29], with ipTM annotated). As shown in Fig. 4a, NACraft generalized across both RNA–protein and DNA–protein contexts under de novo and similarity-guided design, reaching target-wise maximum ipTM values of 0.93 for RNA and 0.91 for DNA, with the maximum ipTM exceeding 0.75 for every target. Fig. 4b further shows that the redesign stage improved candidate quality, particularly for challenging cases such as 8X0N, for which the maximum ipTM increased by nearly 0.3. Consistent improvements were also observed in maximum ipTM, maximum pLDDT and minimum iPAE (Fig. 4c–e). As in the therapeutically relevant protein-target benchmark, ipTM correlated positively with aptamer pLDDT in both de novo design (Pearson *r* = 0.748, *P* = 8.5 × 10^−71^) and similarity-guided design (*r* = 0.774, *P* = 2.5 × 10^−80^; Fig. 4f), supporting agreement between predicted interface formation and aptamer-fold confidence. At the candidate level, the mean fractions with ipTM ≥0.60 were 52.55% for RNA and 67.36% for DNA; under a more stringent ipTM ≥0.70 threshold, a substantial fraction of candidates remained high-confidence (27.11% for RNA and 38.17% for DNA). As shown in Fig. 4g, the target-level candidate-fraction distributions of similarity-guided design across different ipTM thresholds were slightly better than that of de novo design, showing that similarity-guided design generates high-confidence candidates effectively with the guidance of sequence information. Furthermore, we quantified the effect of sequence-sampling number using best-of-*N* analysis. For de novo design, the resampled best ipTM increased from 0.64 at *N* = 1 to 0.79 at *N* = 10, 0.82 at *N* = 100 and 0.855 at *N* = 300. Similarity-guided design increased from 0.63 at *N* = 1 to 0.79 at *N* = 10, 0.84 at *N* = 100 and 0.86 at *N* = 300 (Fig. 4h). The largest gains occurred within the first tens of sampled candidates, followed by progressively smaller improvements at larger sampling numbers. Collectively, these results show that NACraft supports both RNA and DNA aptamer generation, accommodates de novo and similarity-guided settings, and benefits from broader candidate sampling across heterogeneous design contexts.

**Figure 3.**
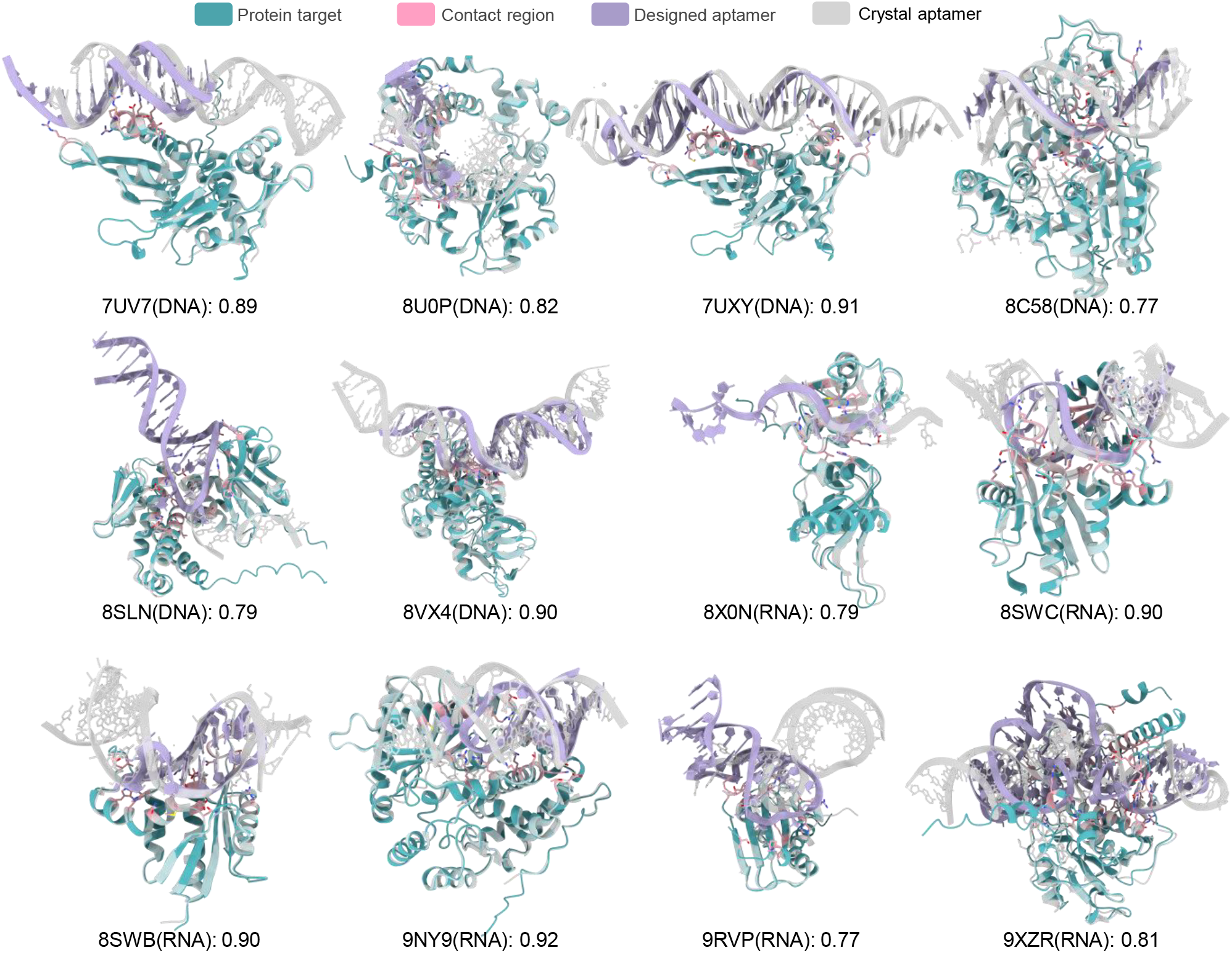
Representative NACraft-designed complexes across the NA-12 benchmark. For each of the six RNA and six DNA targets, the AF3-predicted complex with the highest ipTM is shown.

**Figure 4.**
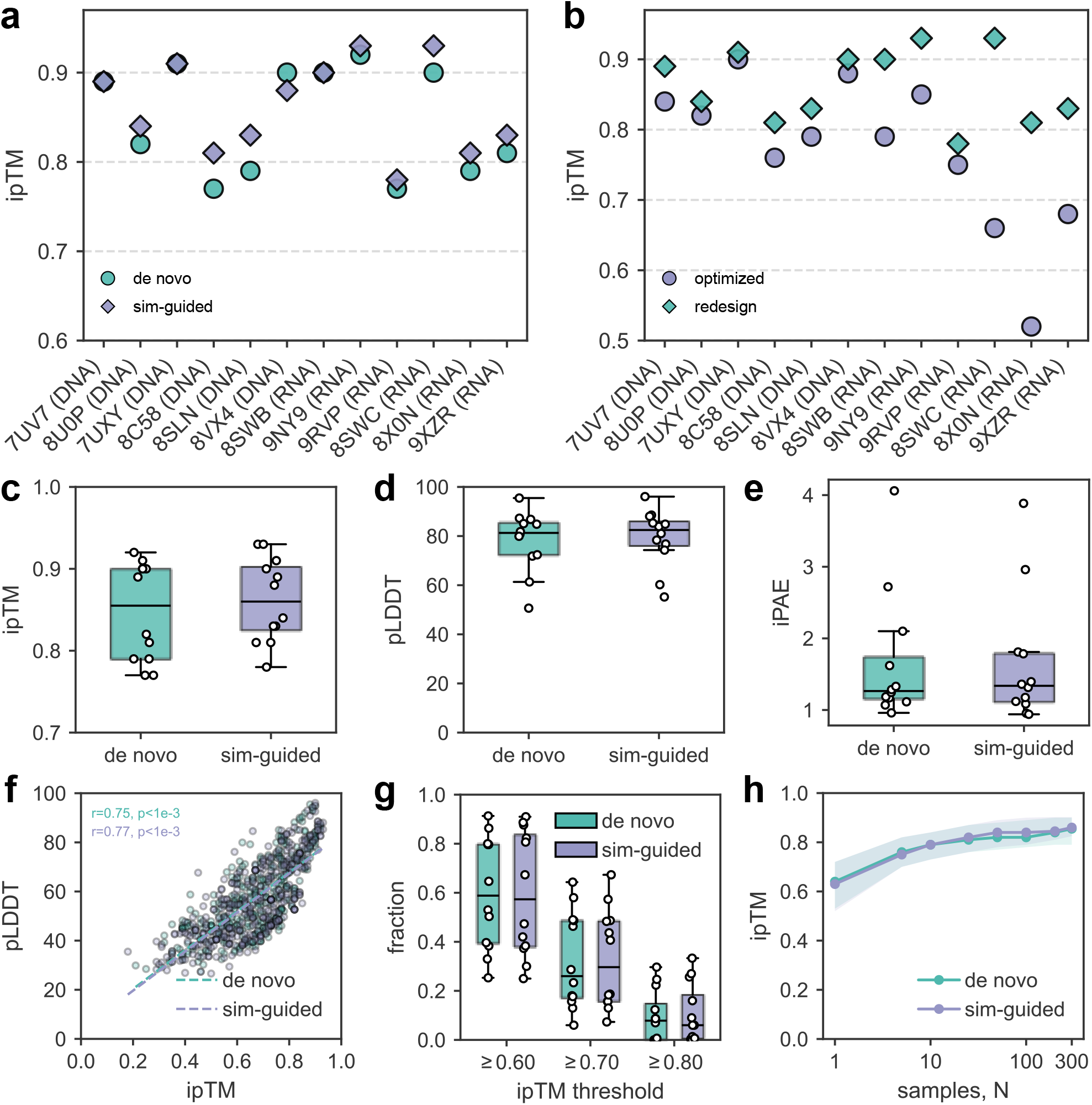
NACraft generated high-confidence RNA and DNA candidates across NA-12. **a**, Target-wise maximum AF3 ipTM for de novo and similarity-guided designs. **b**, Target-wise maximum AF3 ipTM for directly optimized sequences and NA-MPNN-redesigned candidates. **c**, Target-level maximum AF3 ipTM by design mode. **d**, Target-level maximum aptamer pLDDT by design mode. **e**, Target-level minimum AF3 iPAE by design mode. **f**, Candidate-level AF3 ipTM versus aptamer pLDDT for de novo and similarity-guided designs; dashed lines denote linear fits and panel annotations report Pearson correlations. **g**, Distributions of target-level candidate fractions at AF3 ipTM thresholds of *≥* 0.60, *≥* 0.70 and *≥* 0.80, grouped by design mode; boxes show the interquartile range, centre lines denote medians and points denote individual targets. **h**, Resampled best AF3 ipTM as a function of sampling budget *N*; the line denotes the best-of-*N* estimate and shading denotes the interquartile interval across 200 deterministic nested resamples per target and design mode. NA-12 comprises six RNA and six DNA targets and 7,200 AF3-scored candidates.

### 2.4 NACraft enables target-selective aptamer design

Target-selective aptamer design requires discrimination between closely related molecular surfaces, as high affinity toward a desired target does not guarantee avoidance of homologous proteins. We therefore selected the EGFR [25]-HER2 [8] pair as a stringent model of differential molecular recognition. EGFR and HER2 are closely related members of the ErbB receptor family and share substantial structural similarity, making them a challenging test case for evaluating whether NACraft can optimize target preference while reducing recognition of competing surfaces. EGFR domain III was used as the desired target and HER2 domain III as the competing target (Fig. 5a and b; Supplementary Tables 1 and 4). Each candidate was independently evaluated by AF3 against both proteins, allowing selectivity to be measured directly rather than inferred from single-target confidence. In this experiment, NACraft generated 1,800 candidates, of which 69.44% had a higher ipTM for EGFR than for HER2 (Fig. 5c–e). We further defined the differences as ΔipTM = ipTM_EGFR_ − ipTM_HER2_ and ΔiPAE = iPAE_HER2_ − iPAE_EGFR_. As shown in Fig. 5c–e, RNA designs showed stronger predicted selectivity than DNA designs: 82.56% of RNA and 56.33% of DNA designs favoured EGFR by ipTM, while 79.11% and 59.00%, respectively, favoured EGFR by iPAE. Concordance between the two interface metrics therefore supports a target-selective signal, with RNA showing the clearest selectivity in this campaign.

**Figure 5.**
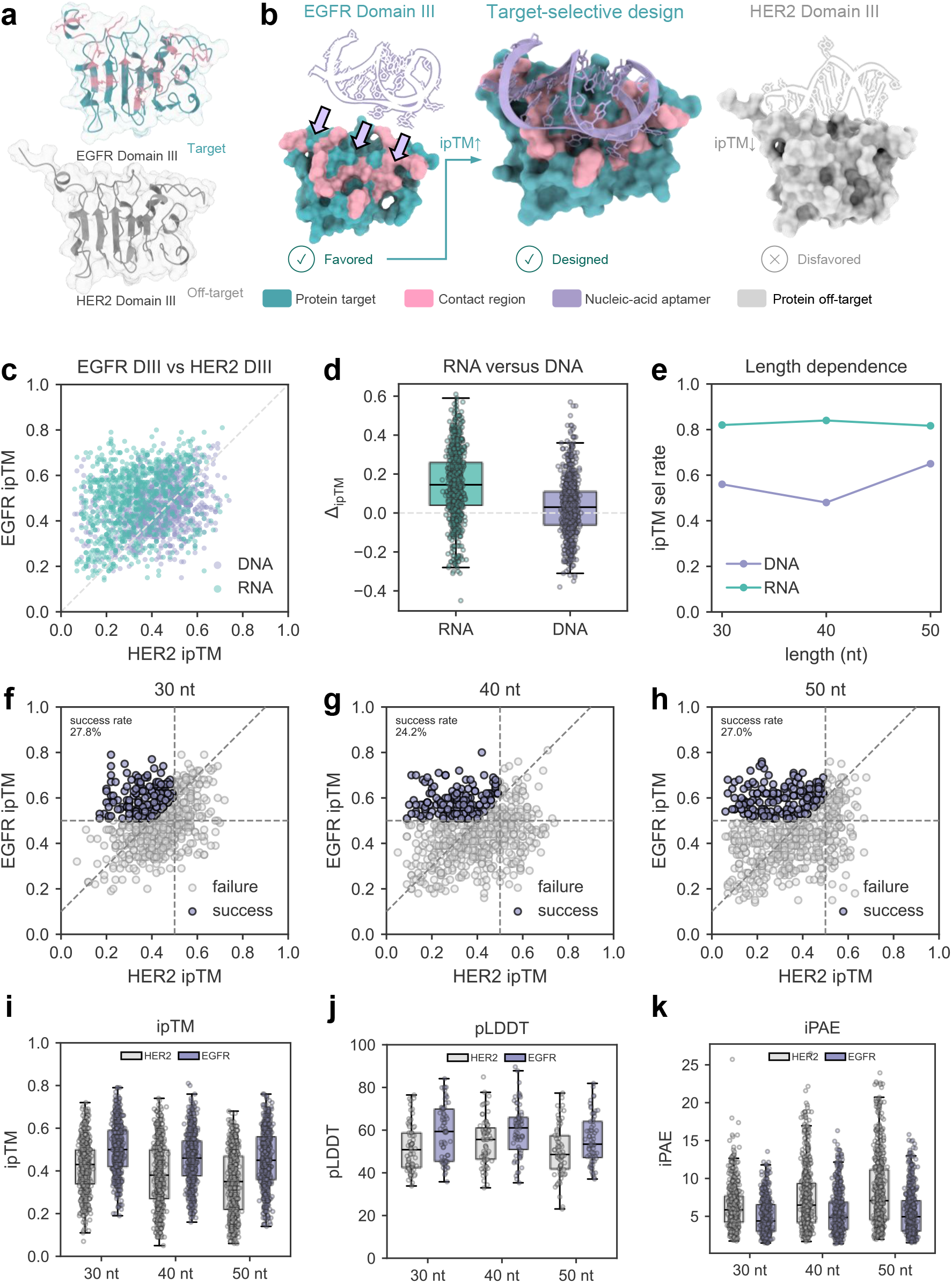
NACraft generated target-selective aptamers between EGFR and HER2 domain III. **a**, Structure visualization of target EGFR domain III and off-target HER2 domain III. **b**, Illustration of designing a nucleic-acid aptamer that favours the intended protein target while disfavouring the off-target protein. **c**, EGFR and HER2 AF3 ipTM values for RNA and DNA designs. **d**, Distribution of ΔipTM = ipTM_EGFR_ *−* ipTM_HER2_. **e**, Length-dependent fraction of candidates with ipTM_EGFR_ *>* ipTM_HER2_. **f--h**, Candidate-level ipTM_HER2_ versus ipTM_EGFR_ for 30, 40, and 50 nt designs. Purple points satisfy all three success criteria: ipTM_HER2_ *<* 0.50, ipTM_EGFR_ *>* 0.50 and ipTM_EGFR_ *>* ipTM_HER2_ + 0.10; grey points fail at least one criterion. **i--k**, Candidate-level AF3 ipTM, aptamer pLDDT and iPAE distributions by aptamer length, grouped by HER2 and EGFR evaluation context. The benchmark comprises 1,800 candidates evaluated against both targets.

**Extended Data Fig. 1.**
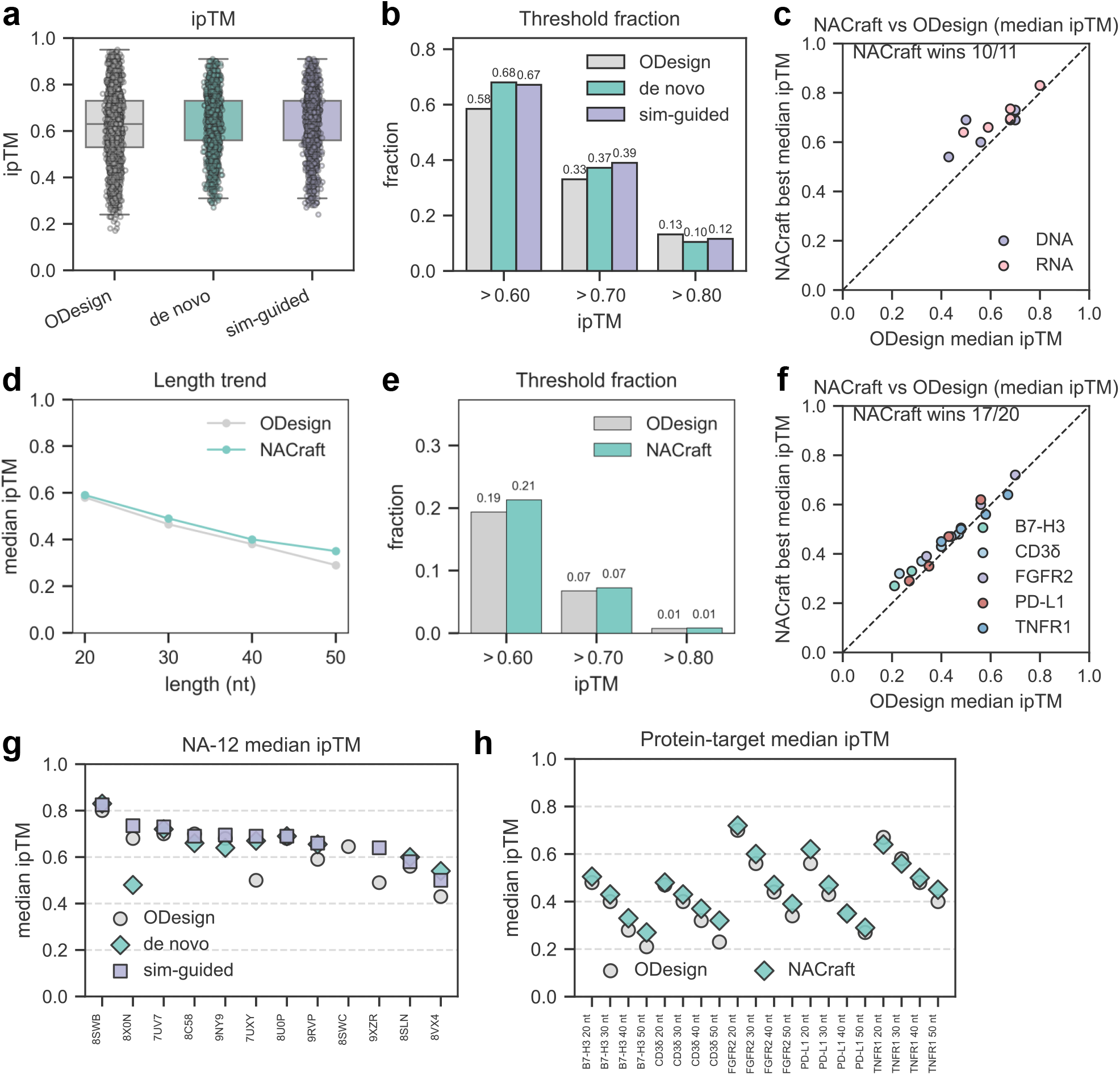
Matched independent AlphaFold3 evaluation of NACraft and ODesign across NA-12 and the protein-target benchmark. **a**, Candidate-level AF3 ipTM distributions for ODesign, NACraft de novo design and NACraft similarity-guided design on NA-12. **b**, Fractions of candidates above AF3 ipTM thresholds of 0.60, 0.70 and 0.80 for the three methods on NA-12. **c**, Paired target-level median AF3 ipTM for ODesign and the better-performing NACraft mode on each canonical-valid NA-12 target; the diagonal denotes equal performance. **d**, Median AF3 ipTM as a function of RNA length for ODesign and NACraft across the five protein targets. **e**, Fractions of candidates above AF3 ipTM thresholds of 0.60, 0.70 and 0.80 for ODesign and NACraft across therapeutically relevant protein targets. **f**, Paired target–length median AF3 ipTM for ODesign and NACraft across the protein target benchmark; the diagonal denotes equal performance. **g**, Target-level median AF3 ipTM for ODesign, NACraft de novo design and NACraft similarity-guided design across NA-12. **h**, Target-level median AF3 ipTM for ODesign and NACraft across B7-H3, PD-L1, CD3*δ*, TNFR1 and FGFR2. The comparison uses AF3 for evaluation. The plotted data comprise 3,400 ODesign, 1,892 NACraft de novo and 1,882 NACraft similarity-guided NA-12 candidates, together with 5,700 ODesign and 6,000 NACraft protein target candidates.

A relative preference for EGFR over HER2 does not by itself establish both high-confidence target binding and effective off-target discrimination. We therefore applied a more stringent joint criterion requiring ipTM_HER2_ *<* 0.50, ipTM_EGFR_ *>* 0.50 and an EGFR-over-HER2 ipTM margin greater than 0.10. Even under this criterion, 27.83%, 24.17% and 27.00% of candidates satisfied all three requirements for 30-, 40- and 50-nucleotide aptamers, respectively (Fig. 5f–h). Beyond threshold-based screening by ipTM, the full metric distributions provided complementary evidence for target selectivity. Across aptamer lengths, candidates showed a consistent shift towards higher ipTM and pLDDT and lower iPAE for EGFR than for HER2 (Fig. 5i–k), indicating greater confidence in EGFR-bound structures and lower predicted interface error, and thus a systematic preference for the positive EGFR target over the competing HER2 off-target. Together, these results show that NACraft can jointly optimize positive- and negative-target objectives during sequence design, extending aptamer generation from single-target binding to target-selective design.

### 2.5 NACraft improves aptamer design over diffusion-based generation

To compare hallucination-based aptamer design with a learned diffusion-based generative approach, we benchmarked NACraft against ODesign [51] under matched AF3 rescoring (Extended Data Fig. 1). We focused on median-based comparisons to assess distribution-level performance.

On the five therapeutically relevant protein targets, NACraft achieved a candidate-level median ipTM of 0.46, compared with 0.43 for ODesign across all length settings (Fig. 1d). This distributional shift persisted at two confidence thresholds: the fractions of candidates above ipTM ≥ 0.60 were 21.33% for NACraft and 19.39% for ODesign, while the fractions above ipTM ≥0.70 were 7.27% and 6.75%, respectively (Fig. 1e). NACraft had a higher median ipTM in 17 of 20 paired target–length settings, with a median ΔipTM = ipTM_NACraft_ − ipTM_ODesign_ of 0.03 (two-sided Wilcoxon signed-rank test, *P* = 8.2 × 10^−4^) (Fig. 1f). In contrast, maximum-ipTM comparisons were less directional: each method achieved the higher maximum in 9 of 20 settings, and two settings were tied. The stronger separation in medians and in the fractions of candidates above both thresholds indicates that NACraft improved the overall candidate distribution rather than relying on isolated best-of-*N* successes. Target-level median ipTM values are shown in Fig. 1h.

The NA-12 benchmark provided a complementary comparison across heterogeneous RNA and DNA contexts. Both NACraft modes reached a candidate-level median ipTM of 0.66, compared with 0.63 for ODesign (Fig. 1a). At ipTM ≥0.60, candidate fractions were 68.02% for NACraft de novo design, 67.16% for similarity-guided design and 58.47% for ODesign; at ipTM ≥0.70, the corresponding fractions were 37.21%, 39.00% and 33.06% (Fig. 1b). When the better NACraft mode was selected independently for each target, NACraft had a higher target-level median ipTM in 10 of 11 paired targets (Fig. 1c). Agreement between candidate-level threshold fractions and paired target-level medians therefore supports a broad improvement across design contexts and shows that the advantage was retained when sequence guidance was introduced.

### 2.6 Ablation study of NACraft framework

**Extended Data Fig. 2.**
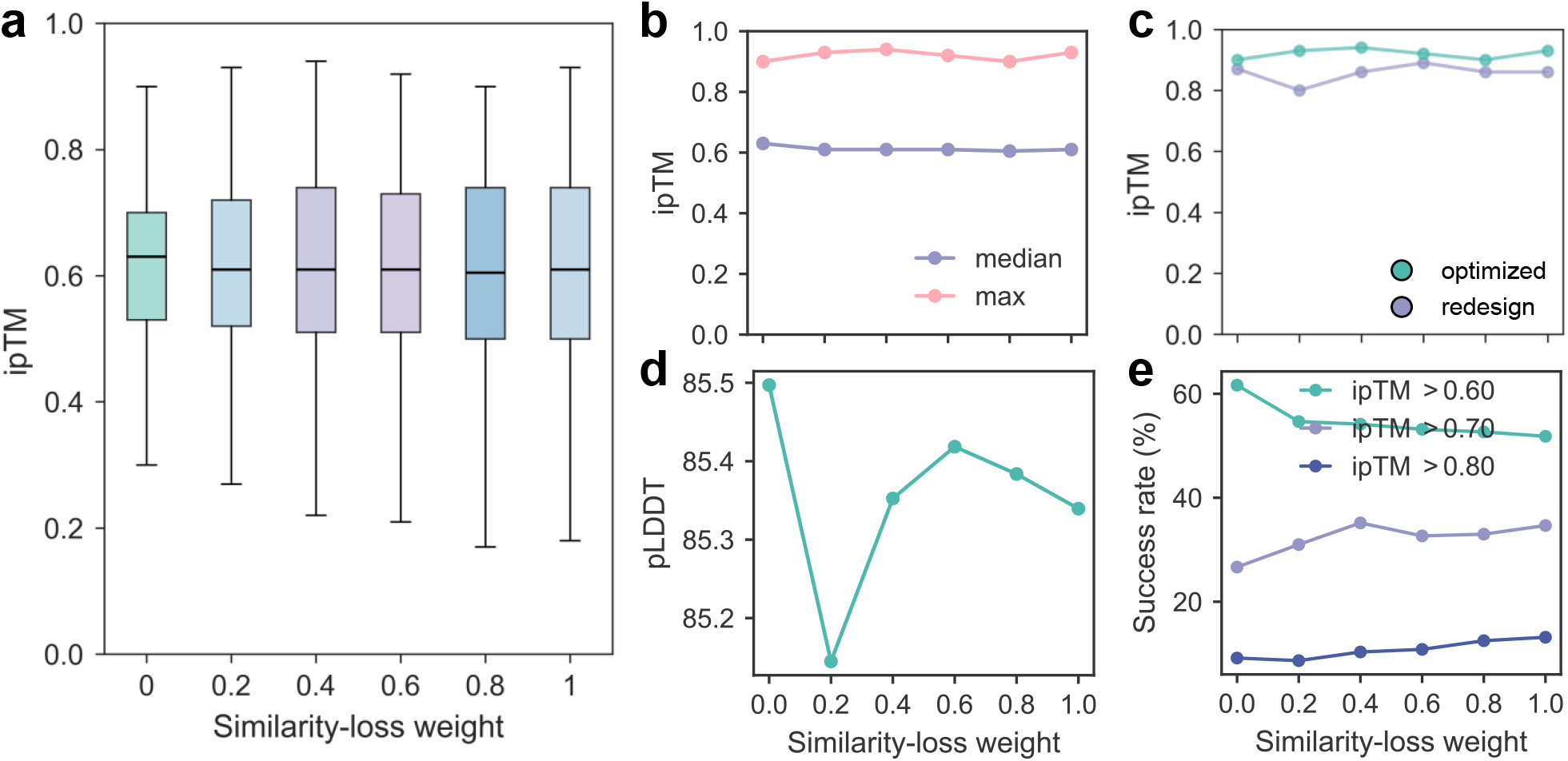
Effect of the sequence-similarity weight on NACraft design. **a**, Candidate-level AF3 ipTM distributions at similarity-loss weights 0, 0.2, 0.4, 0.6, 0.8 and 1.0. **b**, Median and maximum AF3 ipTM at each weight. **c**, Maximum AF3 ipTM for directly optimized and NA-MPNN-redesigned sequences at each weight. **d**, Mean aptamer-chain pLDDT at each weight. **e**, Fraction of candidates above AF3 ipTM thresholds of 0.60, 0.70 and 0.80 at each weight. Each weight comprises 600 AF3-validated candidates from the 12 NA-12 targets, for 3,600 candidates in total.

We next systematically analysed the contributions of key NACraft components, including similarity guidance, sequence redesign and optimization dynamics. The similarity-loss coefficient controls how strongly an input sequence constrains the search. Across weights of 0, 0.2, 0.4, 0.6, 0.8 and 1.0 on NA-12, each represented by 600 AF3-validated candidates, the response was non-monotonic (Extended Data Fig. 2a). A weight of 0.4 produced both the highest maximum ipTM of 0.94 and the highest ipTM ≥0.70 candidate fraction of 35.17%, whereas a weight of 1.0 produced the highest ipTM ≥0.80 candidate fraction of 13.17% (Extended Data Fig. 2b and e). Mean aptamer pLDDT remained stable across all six settings, ranging from 85.14 to 85.50 (Extended Data Fig. 2d). Similarity guidance therefore did not improve monotonically with increasing weight; instead, it controlled the balance between retaining prior sequence information and exploring alternatives, allowing different regions of the high-confidence distribution to be favoured.

**Extended Data Fig. 3.**
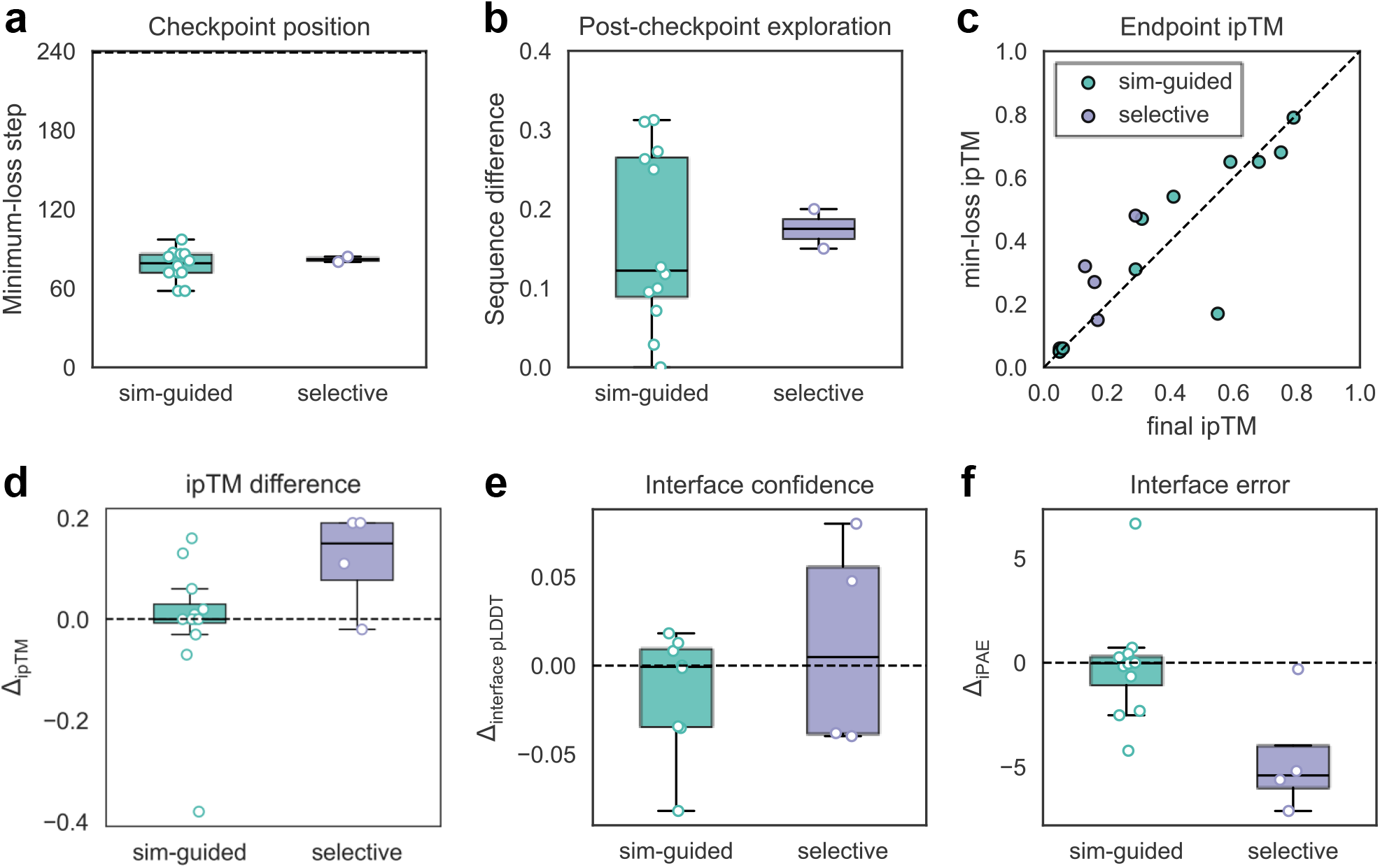
Post-minimum optimization changed sequences while preserving AF3 endpoint quality. **a**, Step at the global minimum total loss; the dashed line marks the final step. **b**, Normalized Hamming distance between minimum-loss and final sequences. **c**, Paired final and minimum-loss AF3 ipTM across 16 context-level evaluations. **d-f**, Paired difference in different metrics including ipTM, pLDDT, and iPAE, where Δmetric = metric_min-loss_ metric_final_. Boxes show medians and quartile ranges; points are context-level evaluations.

NA-MPNN is intended to refine and diversify the sequences produced by differentiable optimization; its contribution can therefore be distinguished from the quality of the initial parent sequences. On NA-12, redesign increased target-wise maximum ipTM for all 12 targets, with the largest gains for the difficult RNA contexts 8SWC (0.66 to 0.93) and 8X0N (0.52 to 0.81) (Fig. 4b). It also increased target-wise maximum aptamer pLDDT for all 12 targets and reduced minimum iPAE for 11 of 12. Analysis of the redesign component across different similarity-loss weights further supported this conclusion (Extended Data Fig. 2c). These concordant changes indicate that NA-MPNN refinement improves both interface confidence and predicted aptamer quality, with the greatest benefits observed in challenging RNA contexts.

The final ablation addressed whether reaching the minimum internal loss marked the end of useful sequence exploration. Optimization-loss trajectories and nucleotide-composition dynamics are reported in Supplementary Figs. 1 and 2, and the retrospective minimum-loss checkpoint analysis is shown in Extended Data Fig. 3. During final annealing, sequences continued to change after the scalar objective reached its global minimum, while independently evaluated AF3 quality remained broadly stable. Thus, post-minimum optimization expanded sequence diversity without a systematic loss of endpoint confidence, although its effect was task-dependent. As shown in Extended Data Fig. 3d–f, similarity-guided design showed no significant difference between minimum-loss and final checkpoints, whereas the target-selective EGFR-over-HER2 task showed better performance at the minimum-loss checkpoint.

## 3 Discussion

Aptamer discovery has historically relied on experimental selection approaches such as SELEX, which iteratively enrich functional sequences from large randomized libraries. Although highly successful, these approaches require extensive experimental screening and provide limited control over the exploration trajectory, target selectivity and rational optimization of sequence–structure relationships. Recent advances in unified all-atom prediction frameworks such as AF3 and Boltz-1 now represent proteins, nucleic acids and their intermolecular interactions within a common structural vocabulary, creating an alternative route towards structure-guided aptamer design. Building on these advances, we developed NACraft, a training-free and programmatic framework for RNA and DNA aptamer design. NACraft optimizes nucleotide sequences through Boltz-1 structure-model feedback and composes molecular contexts with user-defined objective terms to support de novo, similarity-guided and target-selective design within a single formalism. It therefore extends structure-model hallucination from protein binders to nucleic-acid binders and provides a computational complement to selection-driven aptamer discovery without requiring task-specific model training or separate design pipelines.

Across NA-12, NACraft transferred between RNA and DNA while maintaining comparable high-confidence candidate fractions under de novo and similarity-guided design. In the EGFR–HER2 target-selective design task, combining binding and anti-binding objectives shifted candidates towards the intended target, demonstrating that selectivity can be optimized directly to enrich the desired design sequences. Across the therapeutically relevant protein-target benchmark, NACraft generated de novo candidates against structurally diverse surfaces, while matched AF3 evaluation against ODesign showed that its principal advantage lay in shifting candidate distributions rather than producing isolated higher-scoring maxima. Together, these results define NACraft not as a single-purpose aptamer designer, but as an objective-driven framework that can express distinct practical aptamer-engineering problems within the same optimization procedure.

The principal strength of NACraft lies in its programmatic formulation. Rather than defining a fixed generation task, NACraft allows binding, anti-binding, sequence-prior and structural objectives to be recombined according to the intended molecular function. The same optimization engine can therefore search for an unconstrained binder, remodel an experimentally motivated sequence or favour one molecular context over another without retraining the underlying predictor. This flexibility shifts aptamer design from a fixed generation problem towards an objective-driven design process in which target-binding goals can be specified directly during sequence optimization. Component analyses further clarified how this flexibility affects sequence search. NA-MPNN redesign increased the fraction of high-confidence candidates most clearly for RNA, whereas sequence guidance redistributed candidates within the high-confidence tail without producing a monotonic increase in AF3 ipTM. Early-stopped and final candidates achieved broadly similar independently evaluated AF3 quality, but the subsequent annealing and final optimization stages continued to increase sequence diversity without systematically reducing confidence. These later stages may therefore help the search escape local optima and reduce over-specialization to the internal objective, even after the scalar loss has reached its minimum.

Despite the success of NACraft, several limitations remain. First, the reported evaluations rely primarily on AF3-derived confidence metrics and do not directly establish binding affinity, specificity or biological activity. Broader experimental validation will therefore be required to calibrate the relationship between predicted ipTM, pLDDT, iPAE, hotspot contact and measured binding. Second, reliance on a single final verifier may propagate model-specific biases and false-positive predictions. Future implementations could combine AF3, Boltz-family models and Protenix [40] with orthogonal geometric, physics-based or energy-based scoring to prioritize candidates evaluated by multiple independent models. Third, the current implementation is restricted to canonical RNA and DNA nucleotides and does not explicitly model modified nucleotides or an explicit ligand modality. Extending the framework to ligand and protein–ligand contexts would enable the design of nucleic acids against small molecules, ligand-bound proteins and ligand-induced conformational states. Together, these advances suggest a transition from selection-driven aptamer discovery towards programmatic, structure-guided nucleic-acid aptamer design.

## 4 Methods

### 4.1 NACraft design protocol

NACraft designs RNA and DNA aptamers using a four-stage computational protocol. First, a design context specifies the target molecules, nucleic-acid classes, aptamer length and structural objectives. Second, a relaxed nucleotide sequence is optimized by backpropagating differentiable losses through Boltz-1 distogram outputs. Third, selected sequences are structurally materialized and optionally refined and diversified using NA-MPNN. Finally, candidate complexes are independently predicted with AlphaFold 3 (AF3) and filtered using structural- and interface-confidence metrics.

The workflow separates sequence optimization from final structure validation. Boltz-1 provides differentiable geometric feedback during sequence search, with losses evaluated directly on distogram outputs rather than on repeatedly sampled structures. After optimization, Boltz-1 diffusion is used to generate all-atom structures for NA-MPNN redesign and structure-based analyses. AF3 is used only for downstream validation and does not contribute gradients to sequence optimization.

### 4.2 Design-context formulation

NACraft formulates aptamer design as an inverse problem over nucleotide identity. Given a target context *T*, aptamer length *L* and nucleic-acid classes *m* ∈ {RNA, DNA}, the objective is to identify a sequence *S* whose predicted complex satisfies a set of user-defined structural criteria. The design variable is a differentiable distribution over nucleotide identities rather than a discrete sequence or sampled structure.

A design task is specified by one or more evaluation contexts,

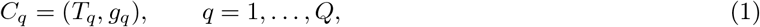

where *T*_*q*_ denotes the molecular context evaluated in state *q*, including the target chain, optional conformer and any additional molecules, and *g*_*q*_ specifies the intended aptamer–context relation, such as contact formation, contact avoidance or similarity to a reference sequence. Single-target design uses one context, whereas target-selective design uses at least one positive and one negative context. The same candidate aptamer sequence is shared across all contexts.

For an aptamer of length *L*, NACraft optimizes the logit tensor

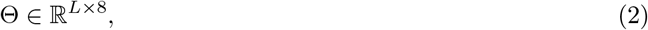

where 8 represents 4 RNA and 4 DNA nucleotides. The `nucleic_acid_class` field defines the valid nucleotide categories A. For RNA,

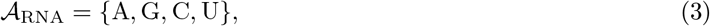

whereas for DNA,

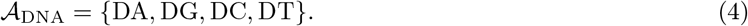

Logits corresponding to invalid tokens are masked by *M*_A_, yielding the relaxed nucleotide distribution

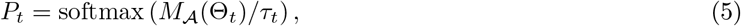

where *τ*_*t*_ is the sequence temperature at optimization step *t*.

The global objective is

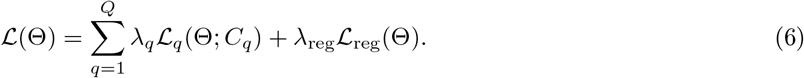

Each structure-dependent objective has the general form

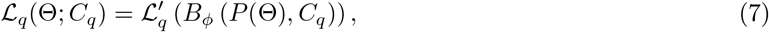

where *B*_*ϕ*_ denotes the frozen Boltz-1 distogram model and *P* (Θ) is the masked relaxed nucleotide distribution. Sequence-based objectives, including similarity to a reference aptamer, act directly on *P* (Θ). This formulation allows geometric constraints, negative-design objectives and sequence priors to be combined within a single optimization problem.

### 4.3 Differentiable sequence optimization

NACraft uses a straight-through estimator to connect discrete nucleotide identities with differentiable optimization. At iteration *t*, the relaxed distribution *P*_*t*_ is projected to the hard nucleotide tensor

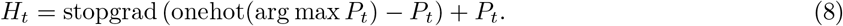

The pseudo-sequence supplied to Boltz-1 is constructed from a scheduled mixture of logits, relaxed probabilities and hard nucleotide assignments. Early iterations therefore explore a continuous sequence space, whereas later iterations increasingly concentrate probability mass on discrete nucleotide identities.

Given the pseudo-sequence and molecular context *C*_*q*_, Boltz-1 predicts pairwise distance distributions

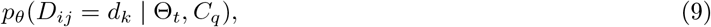

where *D*_*ij*_ is the distance between aptamer token *i* and target or aptamer token *j*, and *d*_*k*_ is the midpoint of the *k*th distance bin. Losses are evaluated directly on these distograms, and the sequence logits are updated according to

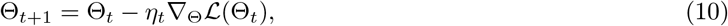

where *η*_*t*_ is the learning rate at iteration *t*.

Following the staged optimization strategy used in BoltzDesign1 [9], NACraft performs 30 warm-up iterations, 100 exploration iterations, 100 annealing iterations and 10 final low-temperature iterations. After each update, gradients corresponding to invalid tokens are set to zero. Fixed motif positions, when specified, are restored before loss evaluation. This procedure constrains optimization to the selected RNA or DNA nucleotides while preserving gradient flow through the Boltz-1 distogram model.

### 4.4 Sequence refinement and diversification with NA-MPNN

Following gradient-based sequence optimization, NA-MPNN is used to refine and diversify selected candidates. This stage expands sequence diversity based on the Boltz-1 backpropagated sequence, whereas the preceding Boltz-guided optimization defines the primary design objective.

Because NA-MPNN requires explicit atomic coordinates, NACraft first generates all-atom structures for each optimized sequence using Boltz-1 diffusion. In the current predictor: boltz configuration, structure generation uses three recycles, 200 sampling steps and five diffusion samples. NACraft writes the resulting complex structures, converts Boltz token identifiers to NA-MPNN nucleotide symbols, performs RNA- or DNA-specific redesign, and converts the redesigned sequences back to standard nucleotide notation. Boltz-1 structures are subsequently regenerated for each redesigned sequence before downstream evaluation. For target-selective tasks, tied redesign is used to ensure that the same aptamer sequence is retained across all positive and negative contexts.

### 4.5 All-atom validation and candidate filtering

All generated candidates are independently evaluated using AF3 in predict-only mode. NACraft writes one AF3 JSON task for each candidate and evaluation context. The designed nucleic acid is assigned to chain A, and target molecules are assigned subsequent chain identifiers. When the target sequence is unchanged, target-specific paired MSAs, unpaired MSAs and template features are computed once and reused across candidates to reduce redundant feature generation.

AF3 outputs are parsed into a common schema containing the model confidence score, interface predicted template modelling score (ipTM), predicted template modelling score (pTM), aptamer-chain predicted local distance difference test score (pLDDT), interface pLDDT where available, and interface predicted alignment error (iPAE). Each candidate is evaluated using five AF3 predictions. Unless otherwise specified, candidate-level summaries use the highest ipTM, highest aptamer-chain pLDDT and lowest iPAE across the five predictions. Aptamer-chain pLDDT is reported on the AF3 scale of 0–100. For the therapeutically relevant target benchmark, normalized interface pLDDT is additionally reported on a scale of 0–1.

Benchmark analyses retain all parse-valid candidates and summarize the complete score distributions. Threshold performance is reported primarily as the fraction of candidates above AF3 ipTM cut-offs of 0.60 and 0.70. Joint filters, such as ipTM *>* 0.60 and aptamer-chain pLDDT *>* 70, are used only for downstream candidate prioritization and do not define the manuscript-wide benchmark population. Protein-aligned nucleic-acid root-mean-square deviation (RMSD) is reported as a retrospective geometric measure.

### 4.6 Objective functions

NACraft adopts a compositional loss interface adapted to nucleic-acid geometry. For RNA and DNA, distance masks are defined using nucleic-acid classes and C1^′^ reference atoms. C1^′^ provides a consistent sugar–base anchor for nucleotide geometry and avoids applying protein-specific C*α* or C*β* conventions to nucleic acids.

The full objective is

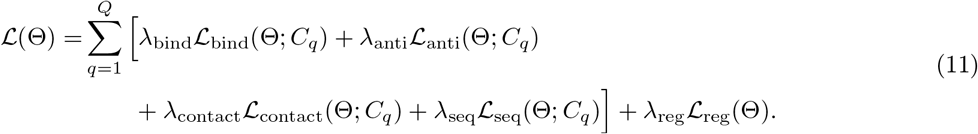

Terms with zero weights are omitted for a given design context, allowing the same implementation to support de novo, similarity-guided and target-selective aptamer design.

#### Binding loss

The binding loss promotes contacts between the aptamer and a target molecule. Let ℬ_*q*_ denote the set of aptamer–target token pairs considered in context *C*_*q*_. For each pair, Boltz-1 predicts a distance distribution over bins with midpoints *d*_*k*_. The probability of contact within distance *d*_0_ is

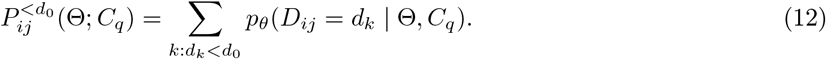

The binding loss is

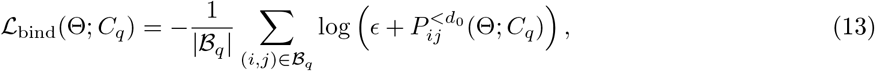

where *ϵ* is a numerical stabilizer. Minimizing this term increases the predicted probability of aptamer–target contacts.

#### Anti-binding loss

The anti-binding loss is applied to an off-target protein or alternative target whose interaction with the candidate aptamer should be disfavoured. It reverses the binding objective:

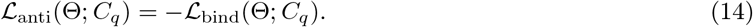

This term is included only for contexts in which target engagement is to be disfavoured.

#### Linear sequence-similarity potential

For similarity-guided design, NACraft uses a reference aptamer sequence

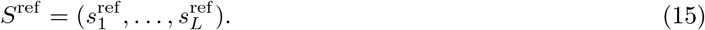

The sequence prior is implemented as a bounded linear potential over the relaxed nucleotide distribution:

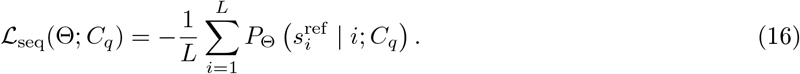

This objective maximizes the mean probability assigned to the reference nucleotide at each position. In contrast to cross-entropy, the linear potential does not impose an increasingly large penalty when the probability of the reference nucleotide is low, thereby retaining flexibility to introduce mutations. Similarity-guided runs combine this potential with reference-based logit initialization. The initialization places the search near the input aptamer, whereas the weighted sequence potential maintains a soft preference for reference identity during interface optimization.

#### Internal-contact loss

The internal-contact loss promotes compact and geometrically plausible aptamer conformations. Let ℐ denote the set of non-local aptamer–aptamer token pairs after excluding neighbouring positions along the sequence. The loss is

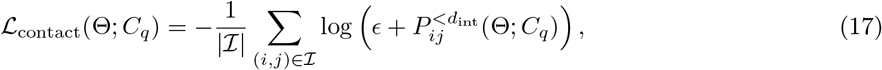

where *d*_int_ is the internal-contact distance threshold. This term is analogous to intramolecular contact objectives used in structure-hallucination methods but is evaluated using nucleotide-specific reference atoms.

#### Regularization loss

The regularization loss stabilizes optimization of the relaxed sequence distribution and can include entropy and logit-norm penalties:

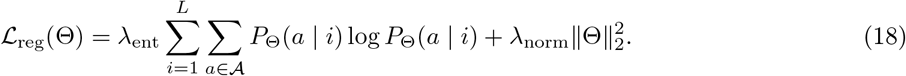

Invalid-token logits are masked before the softmax, and their gradients are set to zero after backpropagation.

### 4.7 Aptamer design modes

NACraft implements three design modes by changing the evaluation contexts and weights assigned to the objective terms. All modes can include binding, internal-contact and regularization losses. Similarity-guided and target-selective design additionally use the sequence-similarity potential and anti-binding loss, respectively.

#### De novo protein-binding aptamer design

For de novo design, nucleotide logits are initialized randomly or uniformly, and no reference aptamer is provided. The active objectives comprise the binding, internal-contact and regularization losses. This mode searches for a previously unspecified RNA or DNA sequence predicted to bind a designated protein target.

#### Similarity-guided aptamer design

For similarity-guided design, nucleotide logits are initialized or biased towards a reference aptamer, and the sequence-similarity potential ℒ_seq_ is added to the objective. The sequence prior is imposed softly, allowing interface optimization to modify positions while retaining information from the input sequence.

#### Target-selective aptamer design

For target-selective design, the same candidate aptamer is optimized against one positive context and at least one negative context. The binding loss is applied to the intended target, whereas the anti-binding loss is applied to an off-target protein or alternative target. Positive and negative contexts therefore contribute directly to sequence optimization, rather than being used only for post hoc filtering.

### 4.8 Benchmark details

#### Therapeutically relevant protein-target benchmark

The de novo benchmark comprised B7-H3, PD-L1, CD3*δ*, TNFR1 and FGFR2. For each protein target, NACraft designed RNA aptamers of 20, 30, 40 and 50 nucleotides using target-specific hotspot residues. Each target–length setting contained 100 directly optimized parent sequences and two RNA-only NA-MPNN children per parent, yielding 6,000 candidates across 20 settings. No reference aptamer sequence was supplied during de novo optimization.

#### NA-12 benchmar

NA-12 comprises 12 protein-binding nucleic-acid targets, including six protein–RNA and six protein–DNA complexes released in the Protein Data Bank after 13 January 2023. RNA targets are within the length intervals of 10–150 nucleotides, while DNA targets are within the length intervals of 10–100 nucleotides. Structures were excluded if they contained ambiguous nucleic-acid sequences, excessively long protein contexts, excessive chain counts or unavailable local coordinate files. Each benchmark entry included the protein structure, nucleic-acid classes, design length, native sequence for retrospective analysis, native protein-interface residues and four default hotspot residues selected from the native interface. For de novo NACraft and ODesign sampling, the native nucleic-acid sequence, backbone and coordinates were withheld from the design algorithms. For retrospective similarity-guided NACraft experiments, the native sequence was supplied only as the sequence prior.

#### AlphaFold3 validation

Benchmark performance was summarized using AF3 ipTM thresholds and, where structural parsing was available, native-patch recovery and RMSD. The primary metrics were AF3 ipTM and aptamer-chain pLDDT; iPAE and protein-aligned nucleic-acid RMSD were treated as secondary metrics. For RMSD calculation, predicted and native protein chains were first superposed using Kabsch alignment. Nucleic-acid RMSD was then computed over the best contiguous shared-length region using available P, C4^′^, C1^′^ or O4^′^ backbone atoms, with unmatched terminal residues excluded when the designed and native nucleic acids differed in length.

#### ODesign

ODesign was run using the odesign_base_na_rigid model, with design_modality set to RNA or DNA and a target-matched sampling budget. ODesign’s internal filtering procedure was disabled, and all sampled sequences were evaluated using the same AF3 rescoring workflow applied to NACraft candidates.

#### Target-selectivity benchmar

The target-selectivity benchmark used a dedicated manifest distinct from the single-target NA-12 benchmark. Each entry specified the nucleic-acid classes, aptamer length, positive target, negative target, source structures, sequences and hotspot annotations. The benchmark evaluated binding strength between EGFR domain III and HER2 domain III, which were treated as homologous but non-identical positive and negative targets, respectively. After sequence generation, each candidate was independently evaluated by AF3 against both targets. The primary readouts were positive-target ipTM, positive-target aptamer pLDDT, negative-target ipTM and the selectivity margin

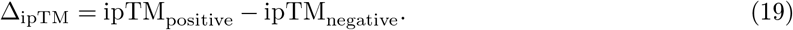

Candidate prioritization required both high positive-target confidence and a positive selectivity margin.

## Data availability

All target structures used in this study were obtained from the Protein Data Bank. Benchmark manifests, processed target structures, analysis data and the source data underlying the reported figures are available through Zenodo [56].

## Code availability

The source code of NACraft is publicly available at Zenodo [56] and GitHub (https://github.com/OTEAM-AI4S/NACraft).

## Author contributions

O.Z. conceived the study and experimental framework. O.Z. and P.A.H. supervised the computational experiments. L.Z. designed and supervised the wet-lab campaign. H.Z. and Jiaqi Wang designed the framework. H.Z. implemented NACraft, performed the computational experiments, analysed the data and drafted the manuscript. W.Z. visualized the structures. Y.X. and Jianmin Wang processed the EGFR and HER2 targets and revised the paper. H.S. carried wet-lab campaign. Q.W., Y.Y. and Z.Y. collected and processed protein targets. G.D. implemented presearch tools. All authors read, contributed to the discussion and approved the final paper.

## Competing interests

The authors declare no competing interests.

## Supplementary information

The supplementary for this paper is publicly available.

## Notes

https://github.com/OTeam-AI4S/NACraft

### Competing Interest Statement

The authors have declared no competing interest.

