## Supplementary for "NACraft: Programmatic nucleic-acid aptamer design via all-atom structure-model feedback"

#### Abstract

Protein–nucleic-acid interactions underpin diverse biological processes and provide a basis for molecular sensing, regulation and therapeutic intervention. However, the coupled dependence of aptamer function on nucleotide sequence, three-dimensional folding and target binding makes rational RNA and DNA binder design challenging. Here we present NACraft, a training-free and programmatic framework for all-atom nucleic-acid aptamer design based on backpropagation through structure-model feedback. By composing binding, sequence-similarity and anti-binding constraints, NACraft supports de novo generation, similarity-guided sampling and target-selective design within a unified optimization framework, without task-specific training or fine-tuning. Computational experiments showed that NACraft generated high-confidence candidates de novo across diverse protein targets, with further improvements achieved through similarity-guided design for both RNA and DNA complexes. Its target-selective design capability was further validated in silico, with 69.44% of paired candidates generated to favour the positive target EGFR over the off-target HER2. Under matched independent AlphaFold3 evaluation, NACraft achieved better performance than ODesign in 10 of 11 NA-12 targets and 17 of 20 protein target–length settings. Together, these results demonstrate the effectiveness and versatility of NACraft and extend structure-model hallucination toward programmatic nucleic-acid aptamer design.

**Code:** `https://github.com/OTEAM-AI4S/NACraft`

### Materials and Methods

#### MSA and template presearch

The MSAs used for Boltz-1 and AF3 folding were precomputed once and cached for subsequent predictions. During candidate evaluation, the cached MSAs were loaded directly to avoid redundant MSA searches and accelerate the folding pipeline. Similarly, template features required by AF3 were precomputed and cached. Both MSA and template features were reused for each target across all generated candidates.

#### Nucleotide-composition dynamics

The annealed relaxation also reshaped nucleotide composition throughout optimization. Continuous stacked-band plots show the fractions of A, C, G and U or T at each optimization step for representative RNA, DNA and target-selective designs (Supplementary Fig. 1a–c), with nucleotide fractions normalized to 100% at each step. Composition continued to change during later optimization stages, including after the scalar objective had plateaued or begun to fluctuate.

#### Optimization-loss trajectories

NACraft optimizes a relaxed nucleotide representation using a differentiable Boltz-1 objective and progressively anneals it towards a discrete sequence. The scalar objective guides the search through sequence space, whereas independent AF3 predictions are used to evaluate the resulting candidates. Across representative optimization trajectories, BindLoss generally decreased during the early stages and subsequently fluctuated as annealing progressively sharpened the nucleotide distribution (Supplementary Figs. 1d–f and **??**). For target-selective design, BindLoss and AntiBindLoss are shown on separate axes because of their different numerical scales. These two terms respectively promote interactions with the desired target context and penalize interactions with the undesired target context.

#### Supplementary Figures

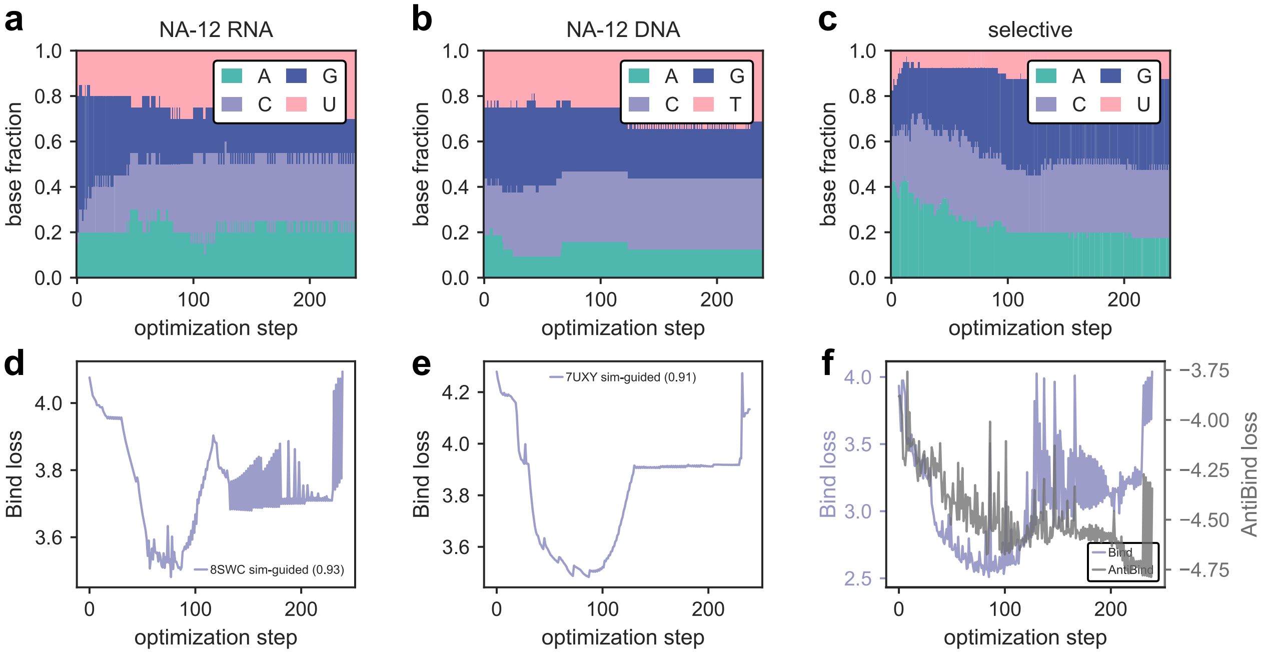


**Figure 1 Annealed optimization continued to reshape nucleotide composition and objective trajectories after their initial descent.** **a**, A/C/G/U fractions across optimization steps for a representative NA-12 RNA design. **b**, A/C/G/T fractions across optimization steps for a representative NA-12 DNA design. **c**, Nucleotide fractions across optimization steps for a representative target-selective design. In **a--c**, nucleotide fractions sum to 100% at each step. **d**, Bind-loss trajectory for the highest-ipTM 8SWC similarity-guided design among candidates with available optimization traces. **e**, Bind-loss trajectory for the highest-ipTM 7UXY similarity-guided design among candidates with available optimization traces. **f**, Bind and AntiBind losses for a representative EGFR–HER2 target-selective design, plotted on separate ordinate axes because of their distinct numerical ranges. The trajectories are representative examples from individual designs.

#### Supplementary Tables

**Supplementary Table 1** Dry-lab design settings and objective coefficients. Parent denotes a directly optimized NACraft sequence and child denotes an RNA- or DNA-only NA-MPNN redesign. All AF3 analyses reused target-specific presearch features.

@p0.17X p0.20p0.27@

| Design block | Context and sampling | Objective coefficients | AF3 analysis |
|---|---|---|---|
| NA-12 de novo | 12 targets (6 RNA and 6 DNA); 100 parents per target; 2 children per parent | $w_{\text{bind}} = 1.0$ | Independent validation |
| NA-12 similarity-guided | Same 12 targets; native/seed sequence initialization; 100 parents per target; 2 children per parent | $w_{\text{bind}} = 1.0$; $w_{\text{sim}} = 0.1$ | Independent validation |
| Target-selective EGFR/HER2 | EGFR DIII positive and HER2 DIII negative targets; RNA and DNA at 30, 40 and 50 nt; 100 parents per context; 2 children per parent | $w_{\text{bind}} = 1.0$ on EGFR; $w_{\text{anti}} = 1.0$ on HER2 | Dual-target validation; success required HER2 ipTM $< 0.50$, EGFR ipTM $> 0.50$ and EGFR ipTM $>$ HER2 ipTM $+0.10$ |
| Protein-target RNA | B7-H3, PD-L1, CD3$\delta$, TNFR1 and FGFR2 at 20, 30, 40 and 50 nt; 100 parents per target–length setting; 2 children per parent | $w_{\text{bind}} = 1.0$ with target hotspots | Independent validation |
| Similarity-weight ablation | 12 NA-12 targets; 10 parents per target and weight; 4 children per parent | $w_{\text{bind}} = 1.0$; $w_{\text{sim}} \in \{0, 0.2, 0.4, 0.6, 0.8, 1.0\}$ | Independent validation |

**Supplementary Table 2 NA-12 target-level AF3 validation metrics.** Each row summarizes 300 AF3-scored candidates for one target and design mode. pLDDT denotes the maximum aptamer-chain pLDDT on the AF3 0–100 scale; iPAE denotes the minimum interface predicted alignment error.

| Target | NA | Length | Mode | $n$ | max ipTM | max pLDDT | min iPAE | ipTM $\geq 0.60$ (%) |
|---|---|---|---|---|---|---|---|---|
| 7UV7 | DNA | 21 | de novo | 300 | 0.89 | 80.75 | 1.62 | 91.3 |
| 7UV7 | DNA | 21 | sim-guided | 300 | 0.89 | 81.04 | 1.81 | 87.7 |
| 7UXY | DNA | 32 | de novo | 300 | 0.91 | 86.80 | 1.17 | 64.7 |
| 7UXY | DNA | 32 | sim-guided | 300 | 0.91 | 88.53 | 1.08 | 67.3 |
| 8C58 | DNA | 14 | de novo | 300 | 0.77 | 50.65 | 4.06 | 79.7 |
| 8C58 | DNA | 14 | sim-guided | 300 | 0.81 | 55.22 | 3.88 | 91.0 |
| 8SLN | DNA | 29 | de novo | 300 | 0.79 | 71.84 | 2.10 | 53.0 |
| 8SLN | DNA | 29 | sim-guided | 300 | 0.83 | 76.77 | 1.78 | 42.0 |
| 8U0P | DNA | 11 | de novo | 300 | 0.82 | 61.34 | 2.72 | 79.7 |
| 8U0P | DNA | 11 | sim-guided | 300 | 0.84 | 60.22 | 2.96 | 82.3 |
| 8VX4 | DNA | 35 | de novo | 300 | 0.90 | 81.80 | 1.11 | 39.7 |
| 8VX4 | DNA | 35 | sim-guided | 300 | 0.88 | 78.35 | 1.18 | 30.0 |
| 8SWB | RNA | 20 | de novo | 300 | 0.90 | 95.47 | 0.96 | 86.3 |
| 8SWB | RNA | 20 | sim-guided | 300 | 0.90 | 96.06 | 0.95 | 88.7 |
| 8SWC | RNA | 20 | de novo | 300 | 0.90 | 84.83 | 1.19 | 38.3 |
| 8SWC | RNA | 20 | sim-guided | 300 | 0.93 | 88.04 | 0.94 | 37.3 |
| 8X0N | RNA | 12 | de novo | 300 | 0.79 | 79.91 | 1.25 | 33.0 |
| 8X0N | RNA | 12 | sim-guided | 300 | 0.81 | 83.89 | 1.12 | 38.3 |
| 9NY9 | RNA | 19 | de novo | 300 | 0.92 | 87.27 | 1.07 | 80.0 |
| 9NY9 | RNA | 19 | sim-guided | 300 | 0.93 | 85.30 | 1.31 | 80.7 |
| 9RVP | RNA | 34 | de novo | 300 | 0.77 | 85.07 | 1.28 | 50.3 |
| 9RVP | RNA | 34 | sim-guided | 300 | 0.78 | 84.85 | 1.36 | 47.3 |
| 9XZR | RNA | 71 | de novo | 300 | 0.81 | 72.38 | 1.33 | 25.3 |
| 9XZR | RNA | 71 | sim-guided | 300 | 0.83 | 74.29 | 1.40 | 25.0 |

**Supplementary Table 3 Therapeutically relevant protein-target AF3 validation metrics.** Each row summarizes 300 RNA aptamer candidates for one target–length setting. pLDDT denotes the maximum normalized interface pLDDT; iPAE denotes the minimum interface predicted alignment error. Hotspot contact is the fraction of candidates with at least one RNA–protein contact within 5 Å of a designated hotspot residue.

| Target | Length | $n$ | max ipTM | max pLDDT | min iPAE | ipTM $\geq$ 0.60 (%) | hotspot contact (%) |
|---|---|---|---|---|---|---|---|
| B7-H3 | 20 | 300 | 0.85 | 0.79 | 1.66 | 25.0 | 40.0 |
| B7-H3 | 30 | 300 | 0.86 | 0.80 | 1.49 | 14.7 | 45.7 |
| B7-H3 | 40 | 300 | 0.73 | 0.76 | 1.70 | 7.7 | 40.0 |
| B7-H3 | 50 | 300 | 0.76 | 0.81 | 1.59 | 2.7 | 35.7 |
| CD3$\delta$ | 20 | 300 | 0.63 | 0.86 | 1.70 | 3.3 | 0.7 |
| CD3$\delta$ | 30 | 300 | 0.59 | 0.87 | 1.63 | 0.0 | 5.0 |
| CD3$\delta$ | 40 | 300 | 0.54 | 0.82 | 1.95 | 0.0 | 3.0 |
| CD3$\delta$ | 50 | 300 | 0.62 | 0.84 | 1.60 | 1.0 | 1.0 |
| FGFR2 | 20 | 300 | 0.88 | 0.84 | 1.26 | 82.3 | 5.0 |
| FGFR2 | 30 | 300 | 0.85 | 0.85 | 1.27 | 51.7 | 6.3 |
| FGFR2 | 40 | 300 | 0.80 | 0.75 | 1.57 | 21.0 | 12.3 |
| FGFR2 | 50 | 300 | 0.76 | 0.75 | 1.63 | 11.0 | 17.0 |
| PD-L1 | 20 | 300 | 0.80 | 0.90 | 1.54 | 60.0 | 32.7 |
| PD-L1 | 30 | 300 | 0.75 | 0.80 | 2.05 | 6.7 | 40.0 |
| PD-L1 | 40 | 300 | 0.63 | 0.78 | 2.00 | 1.3 | 35.0 |
| PD-L1 | 50 | 300 | 0.60 | 0.77 | 2.12 | 0.7 | 26.7 |
| TNFR1 | 20 | 300 | 0.81 | 0.84 | 1.53 | 71.0 | 76.3 |
| TNFR1 | 30 | 300 | 0.75 | 0.80 | 1.66 | 37.0 | 71.0 |
| TNFR1 | 40 | 300 | 0.73 | 0.82 | 1.55 | 19.3 | 73.0 |
| TNFR1 | 50 | 300 | 0.71 | 0.80 | 1.66 | 10.3 | 74.3 |

**Supplementary Table 4 EGFR--HER2 target-selective AF3 validation metrics.** Each row summarizes 300 candidates of one nucleic-acid class and length. EGFR was the positive target and HER2 was the negative target. pLDDT denotes maximum aptamer-chain pLDDT on the AF3 0–100 scale; iPAE denotes minimum interface predicted alignment error. The selectivity fraction requires HER2 ipTM $< 0.50$, EGFR ipTM $> 0.50$ and EGFR ipTM $>$ HER2 ipTM $+0.10$.

| NA | Length | $n$ | EGFR max ipTM | EGFR max pLDDT | EGFR min iPAE | HER2 max ipTM | HER2 max pLDDT | HER2 min iPAE | EGFR $>$ HER2 (%) | selective (%) |
|---|---|---|---|---|---|---|---|---|---|---|
| DNA | 30 | 300 | 0.74 | 79.90 | 1.48 | 0.72 | 76.47 | 1.74 | 56.0 | 18.7 |
| DNA | 40 | 300 | 0.72 | 76.58 | 1.53 | 0.74 | 71.82 | 1.79 | 48.0 | 12.0 |
| DNA | 50 | 300 | 0.69 | 72.57 | 1.58 | 0.68 | 77.37 | 1.97 | 65.0 | 16.7 |
| RNA | 30 | 300 | 0.79 | 84.11 | 1.48 | 0.65 | 74.98 | 1.98 | 82.0 | 37.0 |
| RNA | 40 | 300 | 0.81 | 89.64 | 1.39 | 0.73 | 84.91 | 1.68 | 84.0 | 36.3 |
| RNA | 50 | 300 | 0.76 | 81.91 | 1.51 | 0.67 | 71.62 | 2.04 | 81.7 | 37.3 |
